# Taxonomic and metagenomic analyses define the development of the microbiota in the chick

**DOI:** 10.1101/2022.08.30.505967

**Authors:** Lydia Bogomolnaya, Marissa Talamantes, Joana Rocha, Aravindh Nagarajan, Wenhan Zhu, Luisella Spiga, Maria G. Winter, Kranti Konganti, L. Garry Adams, Sebastian Winter, Helene Andrews-Polymenis

## Abstract

Chicks are ideal to follow the development of the intestinal microbiota and to understand how a pathogen perturbs this developing population. Taxonomic/metagenomic analyses captured the development of the chick microbiota in unperturbed chicks and in chicks infected with *Salmonella enterica* serotype Typhimurium (STm) during development. Taxonomic analysis suggests that colonization by the chicken microbiota takes place in several waves. The cecal microbiota stabilizes at day 12 post-hatch with prominent Gammaproteobacteria and Clostridiales. Introduction of *S*. Typhimurium at day 4 post-hatch disrupted the expected waves of intestinal colonization. Taxonomic and metagenomic shotgun sequencing analyses allowed us to identify species present in uninfected chicks. Untargeted metabolomics suggested different metabolic activities in infected chick microbiota. This analysis, and GS-MS on ingesta confirmed that lactic acid in cecal content coincides with the stable presence of Enterococci in STm infected chicks. Unique metabolites including 2-isopropylmalic acid, an intermediate in the biosynthesis of leucine, was present only in the cecal content of STm infected chicks. Metagenomic data suggested that the microbiota in STm infected chicks contained a higher abundance of genes, from STm itself, involved in branched chain amino acid synthesis. We generated a deletion mutant in *ilvC* (*STM3909*) encoding ketol-acid-reductoisomerase, a gene required for the production of L-isoleucine and L-valine. Δ*ilvC* mutants are disadvantaged for growth during competitive infection with the wild type. Providing the *ilvC* gene *in trans* restored growth of the Δ*ilvC* mutant. Our integrative approach identified biochemical pathways used by STm to establish a colonization niche in the chick intestine during development.

**IMPORTANCE:** Chicks are an ideal model to follow the development of the intestinal microbiota and to understand how a pathogen perturbs this developing population. Using taxonomic and metagenomic analyses we captured the development of the chick microbiota to 19 days post-hatch in unperturbed chicks and in chicks infected with *Salmonella enterica* serotype Typhimurium (STm). We show that normal development of the microbiota takes place in waves, and is altered in the presence of a pathogen. Metagenomics and metabolomics suggested that branched chain amino acid biosynthesis is especially important for *Salmonella* growth in the infected chick intestine. *Salmonella* mutants unable to make L-isoleucine and L-valine colonize the chick intestine poorly. Restoration of the pathway for biosynthesis of these amino acids restored the colonizing ability of *Salmonella*. Integration of multiple analyses allowed us to correctly identify biochemical pathways used by *Salmonella* to establish a niche for colonization in the chick intestine during development.

## INTRODUCTION

Non-typhoidal *Salmonella* (NTS), *Salmonella enterica* subsp. *enterica* serovars Typhimurium and Enteritidis, are the leading cause of bacterial food-borne gastroenteritis in humans and livestock worldwide [1–3]. Since the first report of food poisoning associated with NTS in 1888, infections in livestock and concurrent human cases of food-borne salmonellosis have increased. The overall rate of NTS infections has not declined in 50 years in the United States, and this problem is reflected around the world [4]. The World Health Organization estimates that *S*. Enteritidis and *S*. Typhimurium cause approximately 80% of all human cases, 94 million cases of gastroenteritis worldwide, and 155,000 deaths [3]. Currently in the United States, NTS cause about 1.35 million infections, 26,500 hospitalizations, and 420 deaths annually resulting in $400 million in direct medical costs [5]. There are currently no effective preventative or treatment strategies available_for reducing NTS infection in humans.

NTS gastroenteritis in developed countries is primarily a foodborne infection frequently associated with contaminated chicken meat and eggs [3]. Between 2004 and 2008 approximately 50% of NTS outbreaks were from poultry and eggs (29% and 18% respectively) (CDC National Outbreak Reporting System, 2004-2008). Control methods for NTS in chickens currently include eliminating vertical transmission in breeding stock and replacement chicks, reducing feed and environmental contamination, and improved biosecurity [1]. Infected chickens are difficult to identify because they are sub-clinically colonized with *Salmonella* and have no clinical signs of infection [6]. Broiler chicks contract NTS-infection in the first few days post-hatch and can develop lifelong sub-clinical infection [7, 8]. These subclinical infections with NTS persist in >90% of birds at 8-9 weeks of age (well within the age at which commercial broilers are slaughtered) [8]. The national prevalence of NTS contamination in chicken carcasses is approx. 31% (16.8% *S*. Enteritidis, and 14.5% *S*. Typhimurium) [9].

Reduction in subclinical intestinal carriage of NTS in chickens is a key strategy for reducing human NTS gastroenteritis. Such strategies require understanding of the NTS mechanisms for colonization of the chick intestine to develop new methodologies. Newly hatched chicks have a sterile GI tract and are highly susceptible to deadly NTS infection (< 4 days post-hatch) [10]. In mammals, NTS cleverly exploits the intestinal environment and intestinal inflammation to out compete the intestinal microbiota [11–14]. In mice, NTS employs its Type Three Secretion System-1 (TTSS-1) to promote a massive infiltration of neutrophils that in turn liberate reactive oxygen and nitrogen species [15]. NTS encode multiple oxidases/reductases for terminal electron acceptors and can use these nutrients under anaerobic conditions to catabolize a variety of fermentation products (1,2-propanediol, succinate, ethanolamine, and fructose-asparagine) produced in this inflammatory environment [16]. When infected with S. Typhimurium at 4 days post-hatch, chicks are robustly colonized in the intestinal tract, but do not develop strong heterophilic inflammation [17]. Thus, we expect that some of the mechanisms involved in NTS colonization of the developing chick intestine will be distinct from those of mammals. These new mechanisms will add to the already extensive repertoire of mechanisms utilized by *Salmonella* to dominate the intestinal lumen [17].

To further understand NTS requirements for colonization of the chick GI tract, we used taxonomic, metagenomic, and both untargeted and targeted metabolomic analyses of the microbiota during chick development over the first 19 days post-hatch to capture the development of the microbiota to maturity. Untargeted metabolomics and metagenomics allowed us to identify significant pathways needed by *Salmonella* during colonization of the chick intestine, including branched chain amino acid biosynthesis. Mutational analysis of *ilvC (STM3909*), ketol-acid-reductoisomerase, confirmed the necessity for biosynthesis of L-isoleucine and L-valine by STm during colonization of the chick intestine during early development. Thus, our integrative approach allowed us to identify biochemical pathways used by STm to colonize the developing chick gastrointestinal tract.

## RESULTS

### *Salmonella* infection in chicks does not affect weight gain or induce gross pathological changes

We hatched chicks from SPF eggs and divided them into two groups. One group was left unperturbed, while the other group was infected at 4 days post-hatch with 10^8^ colony forming units (CFU) of *Salmonella enterica* serotype Typhimurium ATCC14028s. We monitored weight gain daily, and STm colonization every two days until 19 days post-hatch. Infected chicks became stably colonized with STm in the ileum, cecum, and colon within five hours post-inoculation and remained so for the duration of our study (Fig 1A). STm infected chicks and uninfected chicks gained weight indistinguishably (Fig. 1B). Furthermore, cecal tissue from days 4, 8 and 19 of age from either uninfected chicks or from chicks infected at 4 days of age, suggests only mild damage to the intestinal epithelium after STm infection (Fig. 1C). This damage was apparent only on day 8 post-hatch (day 4 post-infection) in STm infected chicks, and included mild heterophil infiltration, mild lympho-histiocytic infiltration and mild expansion of the rugae.

**Figure 1.**
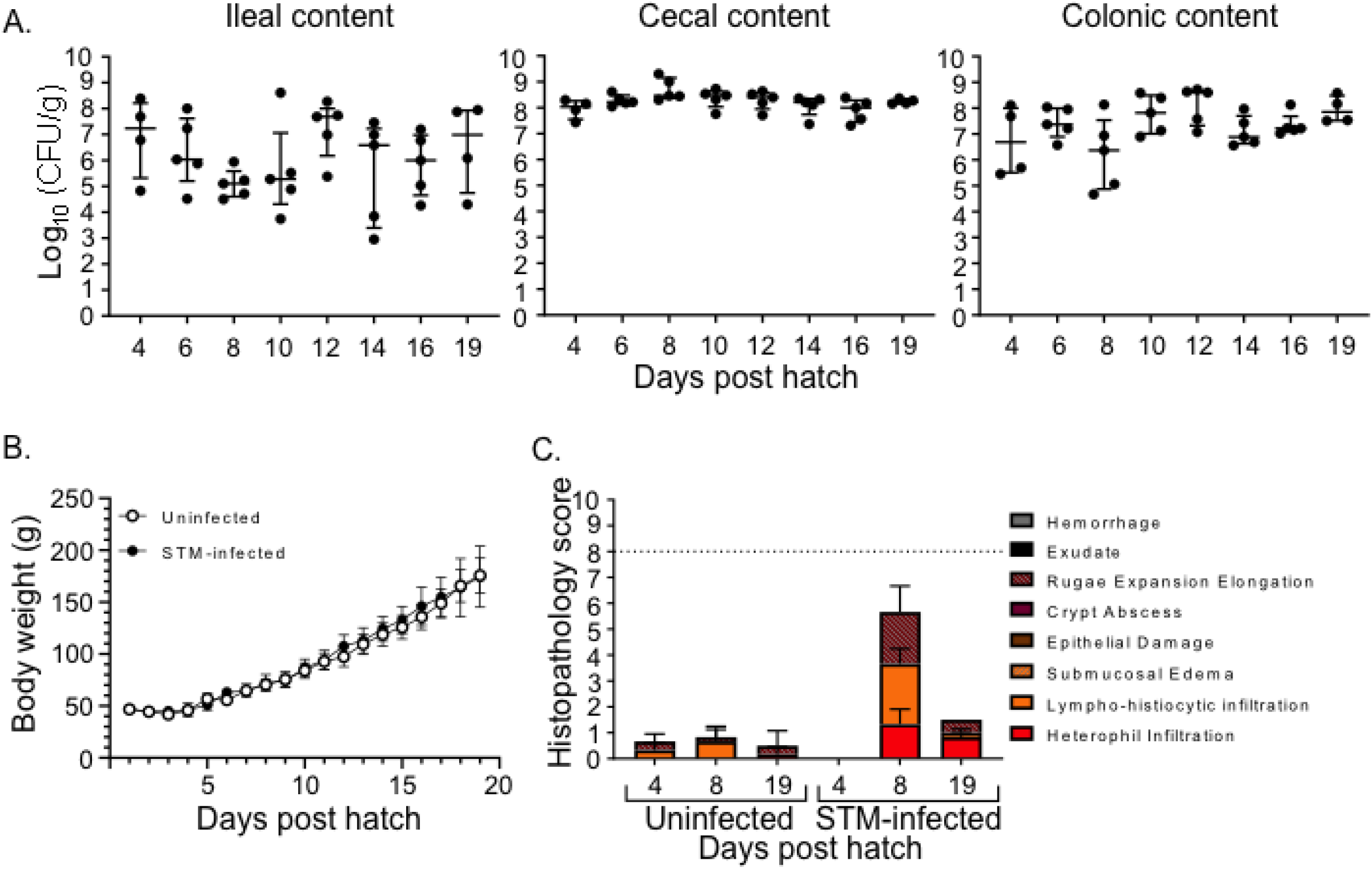
*Salmonella* Typhimurium infection results in prolonged colonization of the intestine without a significant adverse affects on chicks. **A.** Oral inoculation of 4-day-old chicks with 10^8^ CFU *Salmonella* enterica serotype Typhimurium ATCC14028 spontaneously nalidixic acid derivative HA420 leads to stable colonization of ileum, ceca, and colon. **B.** Weight gain of chicks infected with STm compared to uninfected chicks. **C.** Cecal sections were collected from STm infected or uninfected chicks at days 4, 8, and 19 post hatch, fixed in formalin, paraffin-embedded, cut and stained with H&E (Hematoxylin and Eosin). Stained sections were scored for the signs of inflammation. The combined score < 8 corresponds to normal or mild inflammation; the combined score > 8 indicates moderate to severe inflammation

### Cecal microbiota develops in waves and this development is disrupted by STm infection

We sampled the cecal contents of chicks every other day from 2 days after hatching to 19 days of age to define the dynamics of development of the microbiota using 16s rRNA sequencing. Two phyla, Firmicutes and Proteobacteria, dominated the microbiota in developing chicks from hatching to the development of a stable microbiota by day 19 post-hatch, both in unperturbed chicks (Figure 2A) and in chicks infected with STm at day 4 post-hatch (Figure 2C). This finding is in contrast to the intestinal microbiota of mammals, which is more complex and is dominated by Bacteroides and Clostridia [18, 19]. Stability and maturity in the composition of the cecal microbiota at the phylum level occurs between days 10-12 post-hatch.

**Figure 2.**
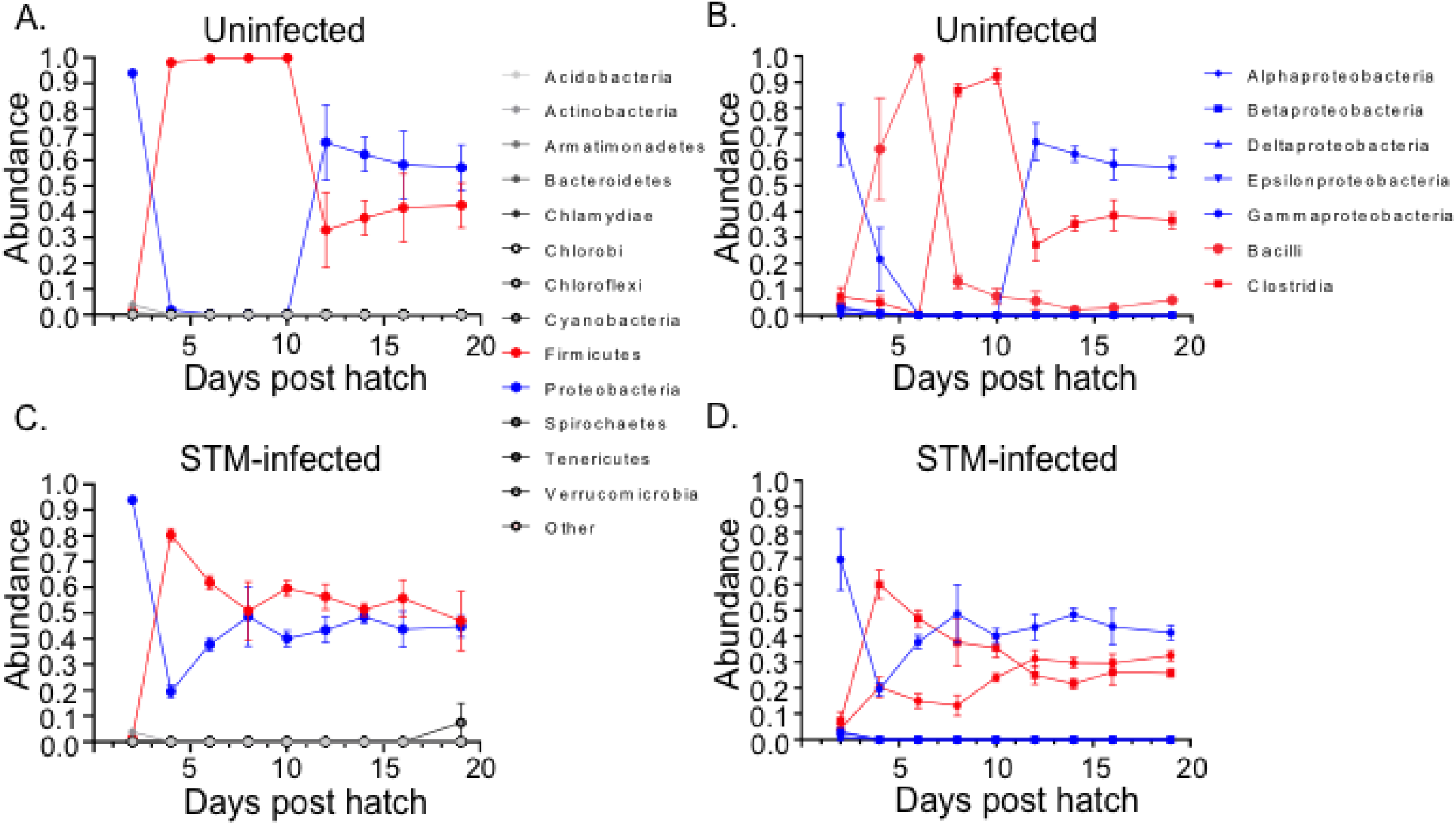
16S rRNA sequencing analysis of chicken cecal microbiota development over first 19 days of life. **A,C.** Composition of microbiota present in the ceca of uninfected (A) or STM-infected (C) chicks at the phylum level. **B,D.** Composition of microbiota present in the ceca of uninfected (B) or STM-infected (D) chicks at the class level.

Looking at the microbiota colonizing the chick ceca at the class level, the ceca of unperturbed chicks are colonized in multiple waves, reaching a stable state by day 12 post-hatch (Fig. 2B). Although sampling at day 2 post-hatch is difficult and the cecal microbiota are sparse, Gammaproteobacteria were the most abundant member of the chick cecal population. The Gammaproteobacteria population diminished by day 6 post-hatch, and Bacilli dominated the microbiota reaching a peak at 6 days post infection. Between days 6 and 8 post-hatch, Bacilli decreased in abundance, and Clostridia dominated the cecal population by day 10 post-hatch. By day 12 post-hatch, Gammaproteobacteria emerged again as approximately 60% of the cecal population, and were followed in prominence by the Clostridiales and to a lesser extent, Bacilli. After day 12 post-infection, the balance of Gammaproteobacteria, Clostridia and Bacilli appeared to be relatively stably maintained until the end of sampling at day 19 post-hatch.

When chicks were orally infected with 10^8^ STm at 4 days post-hatch, STm quickly altered the dynamics of the developing microbiota in the chick cecum (Fig 2D). With the reduction in the relative abundance of the Gammaproteobacteria, the second wave of colonization is by Clostridia, which dominate the population by 8 days post-hatch (day 4 post-infection). At that time, Gammaproteobacteria and Bacilli each represent about 20% of the cecal population while the Clostridia are present in 40-60% relative abundance. By day 8 post-hatch, the Gammaproteobacteria once again are dominant, not surprisingly because these are primarily STm, while the relative abundance of the Clostridia continues to decline. By day 12 post-hatch, Gammaproteobacteria remain dominant, but are closely followed in relative abundance by Bacilli and Clostrida. Thus, quickly after the introduction of STm into the intestinal tract of the developing chick, STm modulates the composition of the cecal microbiota and forms a stable community that differs from that in unperturbed chicks.

### Metagenomic shotgun sequencing identifies species present in developing chick microbiota

In order to identify the species present in the cecal microbiota at different time points post-infection, we performed shotgun sequencing on cecal samples from uninfected and STm infected chicks at days 4, 6, 10, 12, and 19 post-hatch (Fig 3). These experiments confirmed that the most prominent members of the microbiota in unperturbed chicks colonize the ceca in waves. In the first wave of cecal colonization at 4 days post infection, Enterococcus faecalis (Bacilli) is the most highly abundant member of the cecal microbiota. By day 6 post-infection, these appear to be replaced by Enterococcus faecium (Bacilli), and replaced again by day 12 post-infection with Enterobacter cloacae (Gammaproteobacteria). In STm infected chicks, this analysis identified the presence of Bacilli (Enterococcus ssp), and Gammaproteobacteria (*Salmonella*) as expected.

**Figure 3.**
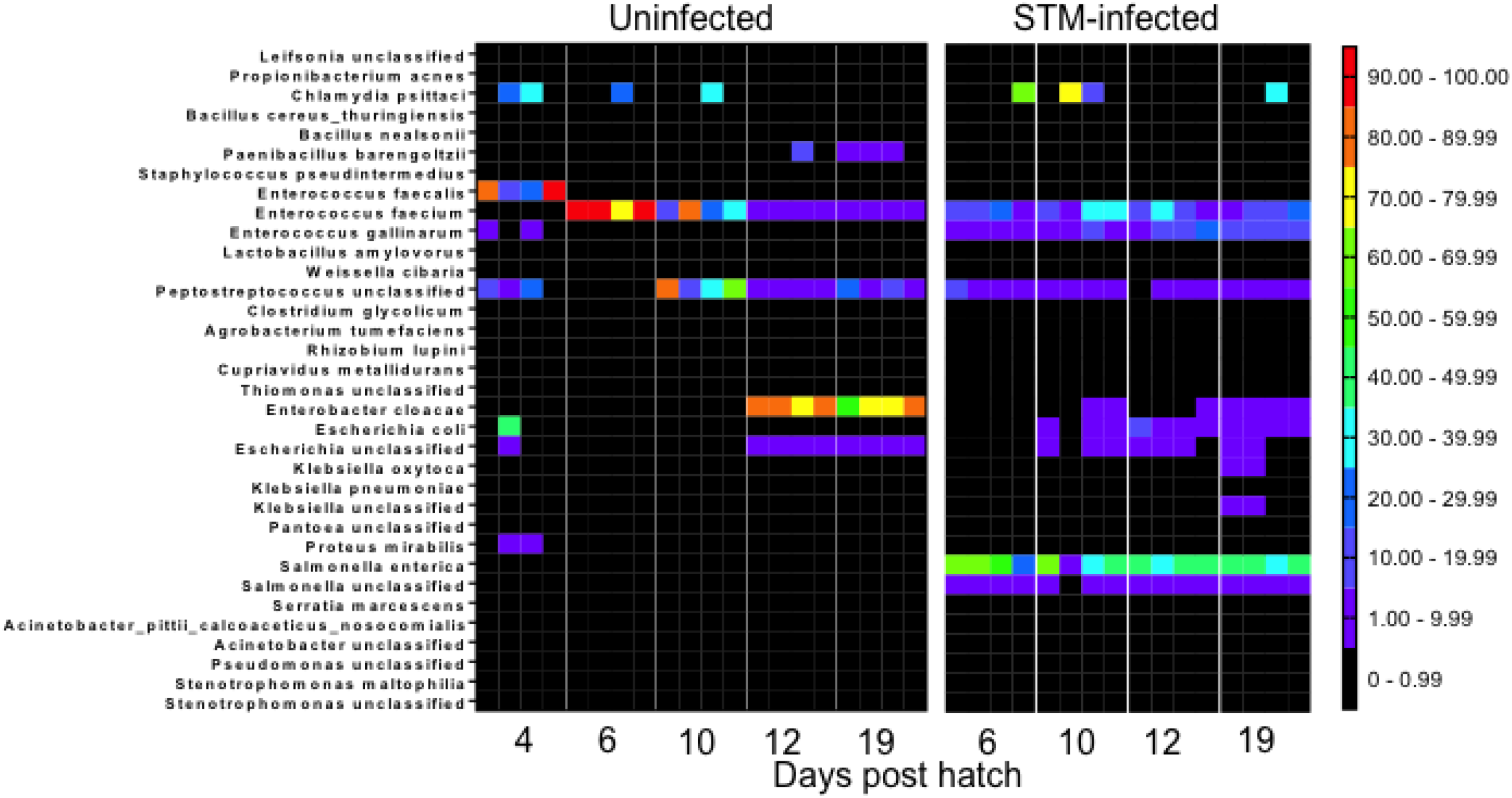
Taxonomic profile of cecal microbiota development using shotgun metagenomic analysis. DNA extracted from the cecal content of uninfected (left panel) or STm-infected (right panel) was used for shotgun sequencing. Taxonomic profiling was analyzed using MetaPhlAn2 and StrainPhlAn2 tools.

### Untargeted metabolomics analysis of cecal contents suggests the presence of common and unique metabolites

We performed an untargeted analysis of metabolites from the ceca of both groups of chick from days 4, 6, 8, and 12 post-hatch (Fig S1), to understand the metabolic activities of both microbial communities. The results of this analysis showed differences in metabolic composition between the unperturbed and STm infected groups (Fig S1). Lactic acid, for example, was detected in higher quantity in the ceca of uninfected chicks versus unperturbed chicks in this analysis. Using GS-MS, we quantified the amount of lactate in the ceca of birds of both groups over 19 days of life (Fig 4). This analysis confirmed the metabolomics result that lactate is present in much higher amount in the ceca of STm infected chicks vs. unperturbed chicks. Finally, this data supports our metagenomic result suggesting that enterococci are an important and stable member of the cecal community in chicks after STm infection.

**Figure 4.**
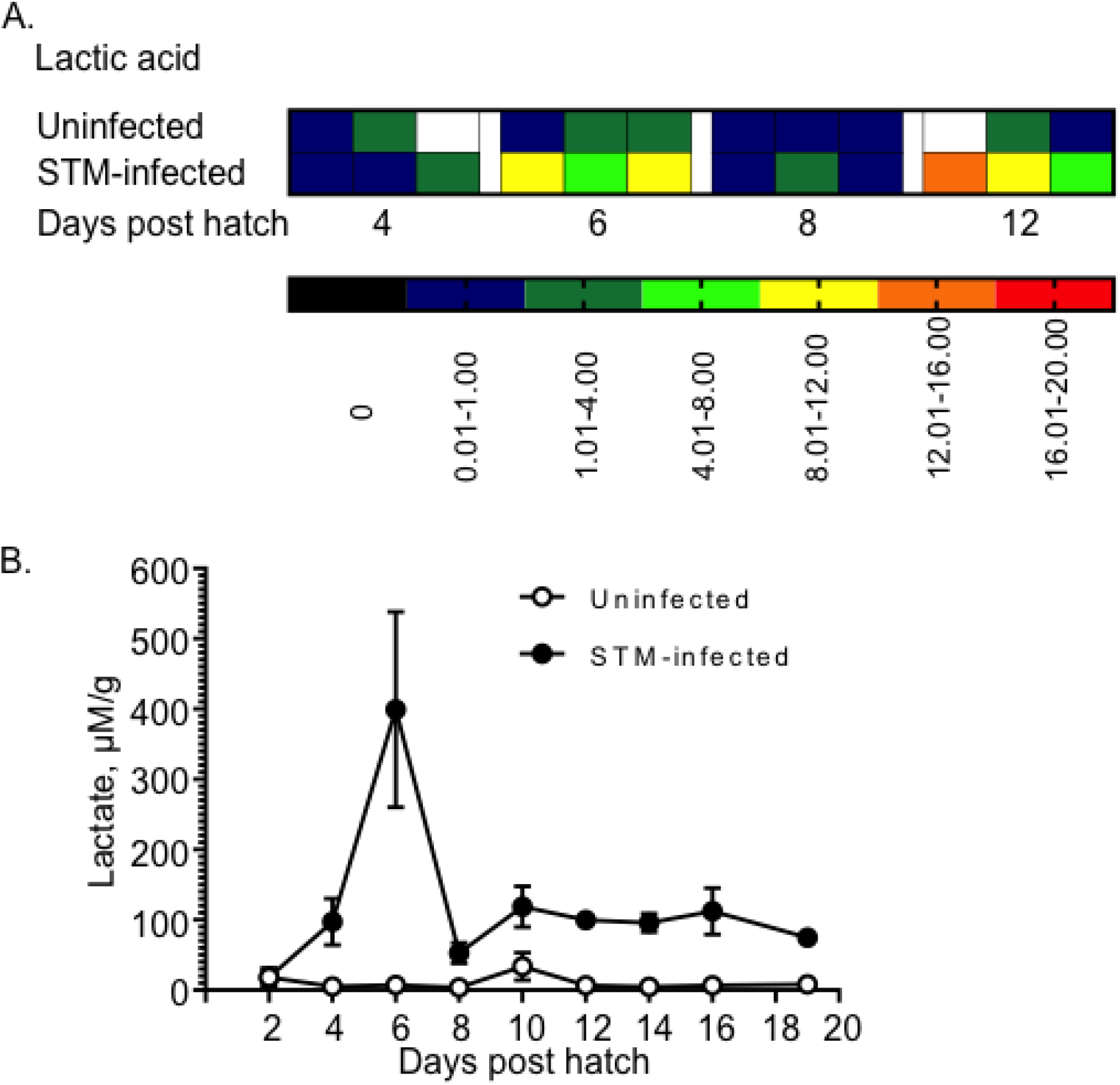
The presence of elevated levels of lactate in the ceca of *Salmonella*-infected chicks compared to uninfected birds was confirmed by GC-MS analysis. **A.** The non-targeted metabolomic approach identified differences in lactic acid concentrations between infected and uninfected chicks (also Figure 4, red box). **B.** Quantification of lactate content in the cecal content of uninfected (open circles) and STM-infected (black circles) chicks using GC-MS multiple reaction monitoring mode.

In addition, metabolites that were present in one group of chicks only were revealed. One of these metabolites, 2-Isopropylmalic acid, was uniquely present in the ceca of chicks infected with STm (Fig S1, Fig 5). 2-Isopropylmalic acid is an intermediate product in the biosynthesis of leucine from 2-ketovaline. Functional profiling of the shotgun sequencing data suggested that the microbial community from STm-infected chicks was characterized by a higher abundance of genes involved in branched chain amino acid biosynthesis (Fig 5). Genes in pathways for the biosynthesis of branched chain amino acids isoleucine and valine were particularly abundant in the ceca of chicks from the day of infection until the termination of the experiment in these two groups, but not in unperturbed chicks. Detailed analysis of this data suggested that the majority of genes contributing to branched chain amino acid biosynthesis belonged to *Salmonella* enterica (Fig 6). These data in combination with the presence of abundant 2-Isopropylmalic acid in STm infected chicks suggested that the biosynthesis of isoleucine and valine are particularly important for STm growth in the developing chick intestine.

**Figure 5.**
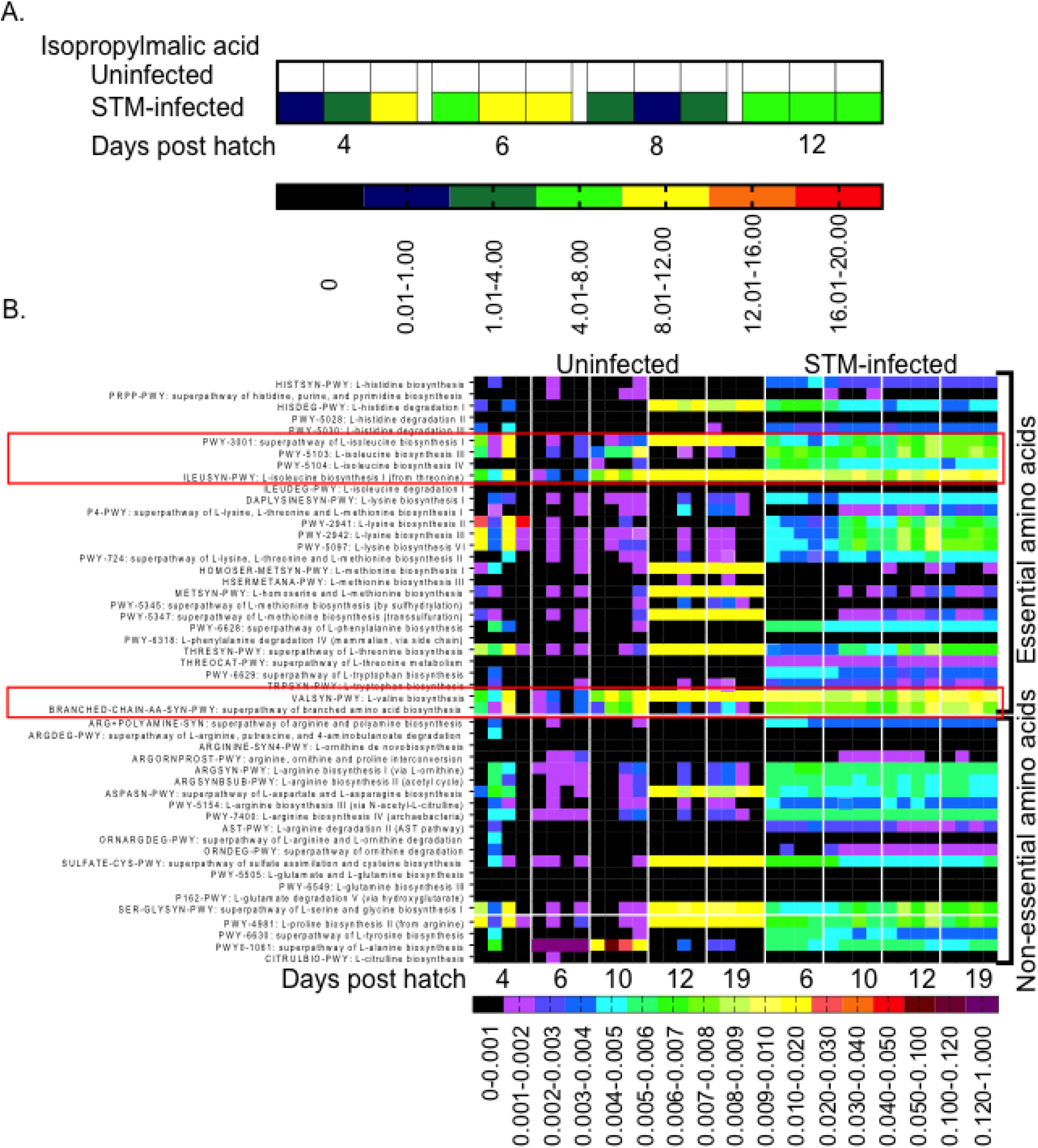
The presence of isopropylmalic acid detected by non-targeted metabolomics, and the higher abundance of genes involved in branched amino acid biosynthesis in the cecal content of STm-infected chicks suggests the association of this pathway with STm colonization. **A.** Isopropylmalic acid, an intermediate product in the biosynthesis of leucine, was detected by untargeted metabolomics only in the cecal samples collected from STm infected chicks (also see Figure 4, red box). **B.** Functional profiling and relative abundance levels of the amino acid biosynthesis and degradation genes present in the ceca of uninfected (left panel) and STM-infected of infected chicks (right panel) was constructed using HUMAnN2 tool.

**Figure 6.**
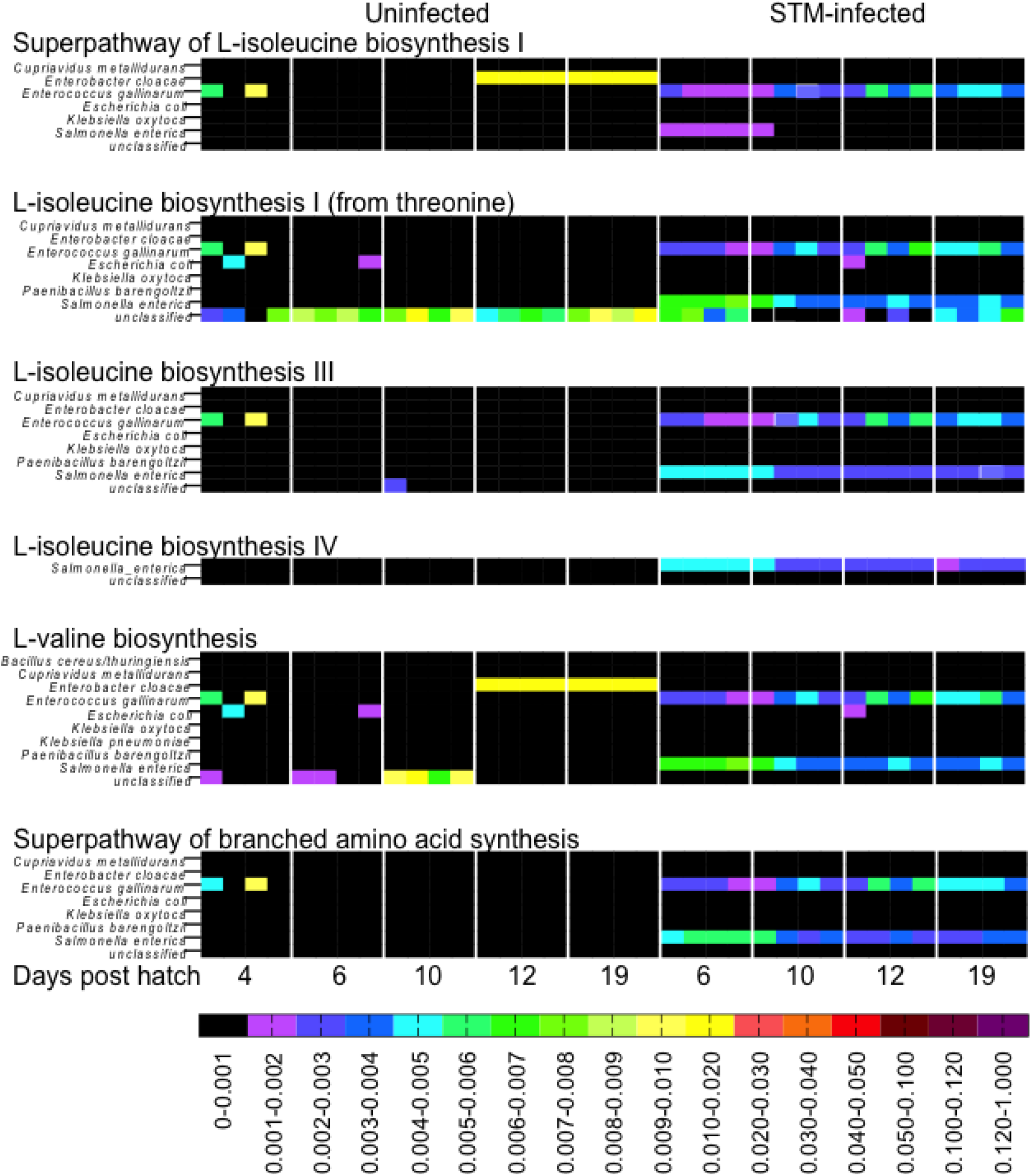
Functional profiling of the shotgun sequencing data indicates that most genes involved in branched amino acid synthetic pathways are from *Salmonella enterica* and *Enterococcus gallinarum* in the infected ceca. The analysis of amino acid biosynthesis and degradation genes present in the ceca of uninfected (left panel) and STM-infected (right panel) of infected chicks was performed using HUMAnN2 tool.

### Branched chain amino acid biosynthesis is critical to STm colonization in the ceca of developing chicks

Both the biosynthesis of L-isoleucine from threonine, and valine from pyruvate require the conversion of (S)-2-acetolactate to (2R)-2/3-dihydroxy-3-methylbutanoate using the enzyme IlvC ((2R)-2,3-dihydroxy-3-methylbutanoate:NADP+ oxidoreductase) (Fig 7A). We generated a deletion mutant in *ilvC* (ΔilvC or ΔSTM3909), and tested the ability of this mutant to grow on minimal media, and minimal media supplemented with L-isoleucine, L-valine, or both. Mutants lacking *ilvC* failed to grow on minimal media, or on minimal media supplemented with either L-isoleucine or L-valine (Fig 7B). Supplementation of minimal media with both L-isoleucine and L-valine or return of an intact *ilvC* gene *in trans* restored the ability of the *ilvC* mutant to grow. Thus, deletion of ilvC creates L-isoleucine and L-valine auxotrophy that is reversible with the introduction of an intact copy of *ilvC* in trans (Fig 7B).

**Figure 7.**
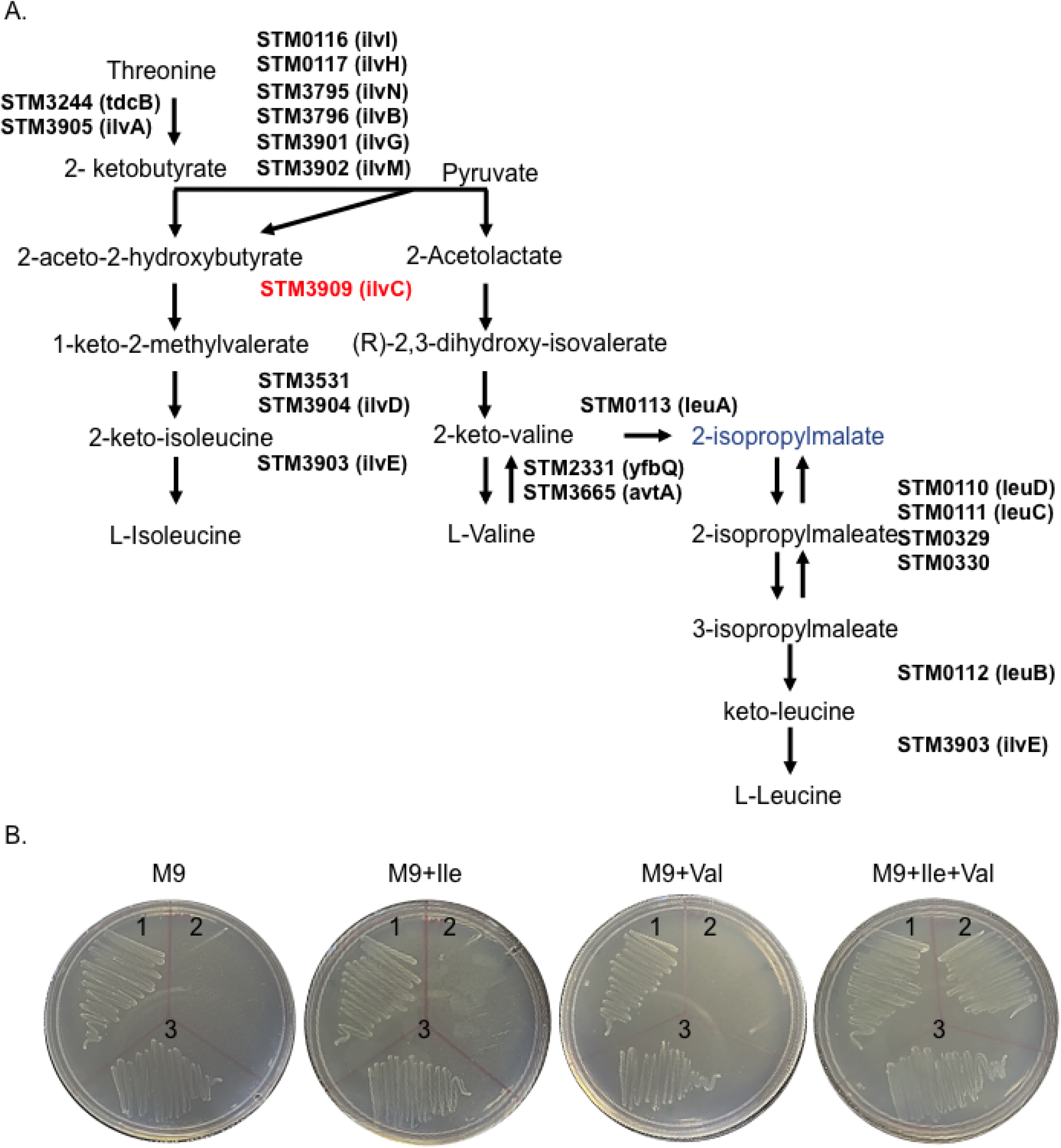
*STM3909 (ilvC*) gene is essential for branched-chain amino acids (BCAA) synthesis in *Salmonella* Typhimurium. **A.** An overview of branched-chain amino acids synthesis in *Salmonella* Typhimurium. **B.** The *STM3909* mutant requires the presence of L-Isoleucine (Ile) and L-Valine (Val) for growth in minimal medium. 1-wild type; 2 - Δ*STM3909 (ilvC*); 3 - Δ*STM3909* pWSK29-*STM3909*.

We tested the ability of a deletion mutant in Δ*ilvC (ΔSTM3909*) to colonize 4-day old chicks using competitive infections with the otherwise isogenic wild type (Figure 8). Although initial colonization between the deletion mutant and the wild type was equal, deletion of *ilvC* severely restricted growth of the mutant in the intestinal tract and systemic sites in chicks during 15 days of infection. Returning an intact copy of *ilvC in trans* restored the colonizing ability of the *ilvC* deletion mutant (Fig 8).

**Figure 8.**
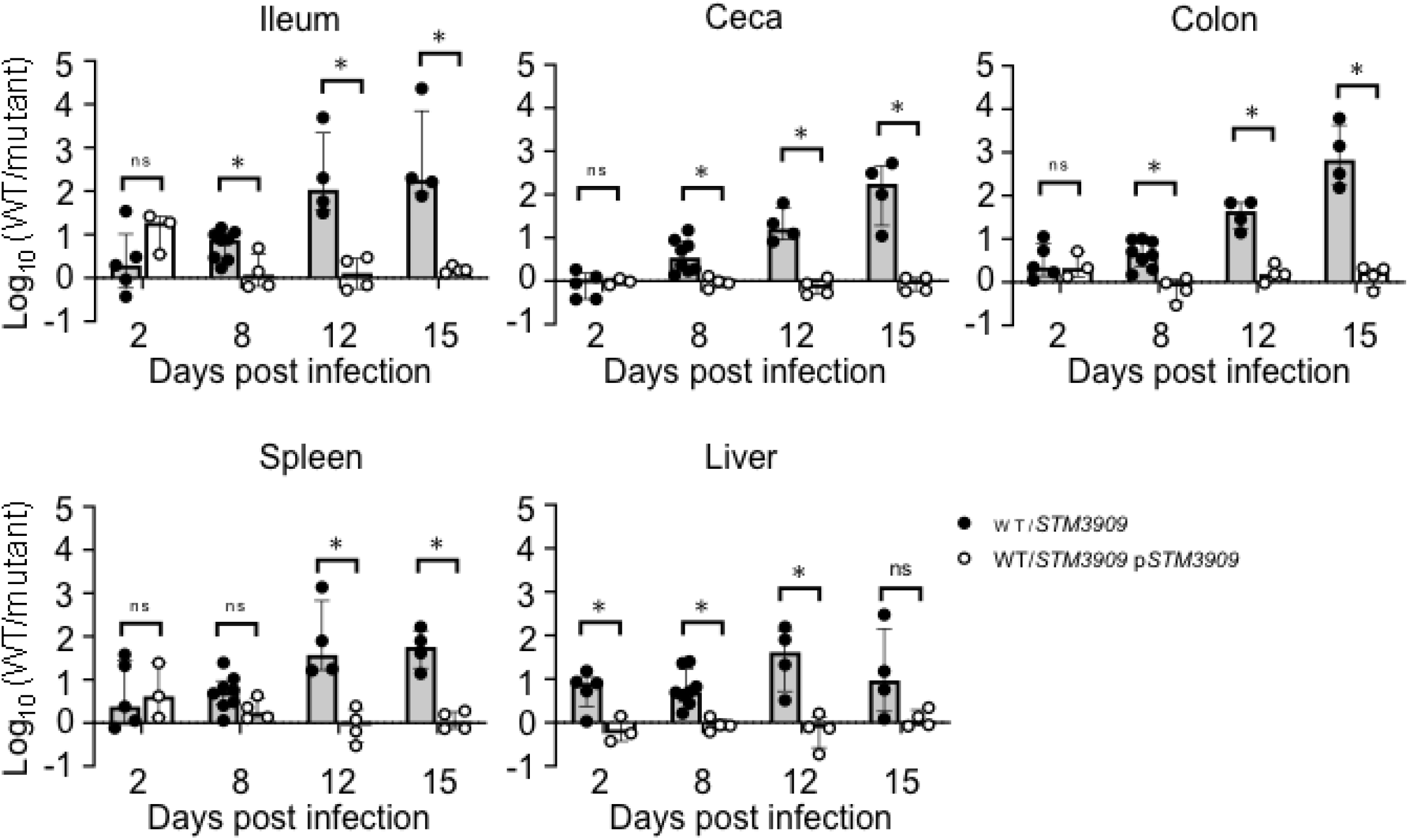
Branched amino acid biosynthesis is required for *Salmonella* colonization of 4-day-old chicks. Chicks were infected with 10^8^ CFU of a 1:1 mixture of ΔSTM3909 mutant and wild type (black circles), or ΔSTM3909 pWSK29-STM3909 mutant and wild type (open circles) strains. Chicks were euthanized on days 2-, 8-, 12-, and 15-days post infection and ileum, ceca, colon, spleen and liver were collected for enumeration of CFU. Statistical significance was determined using a Student’s *t* test. *, significant difference (*P* < 0.05) in the ratio of the mutant/wild type in the collected tissue compared with that of the inoculum.

## DISCUSSION

In modern commercial poultry production, newly hatched chicks have no contact with adult birds, and the microbial communities present in the environment function as the inoculum that can shape the development of chicken gut microbiota. The process of intestinal colonization by bacterial organisms is rapid and very robust. Within one day of hatching, bacterial densities in the ileum and cecum of broiler chickens reach 10^8^ and 10^10^ CFU per gram of ingesta, respectively. Bacterial concentration further increases during first three days of the life of the chick, reaches 10^9^ and 10^11^ CFU per gram of ileal and cecal content, and remains relatively stable for the following 30 days.[20]

In agreement with previous reports [21–23], we found that during the first several days post hatch, the diversity of cecal microbiota is very low and composed primarily of *Proteobacteria*. Exposure of the chicken gut to Enterobacteriaceae at the day of hatch can induce mild, nearly undetectable inflammation leading to immune tolerance of related bacteria including *Salmonella* [24–26]. Accordingly, the ceca of chickens infected with STm are only mildly inflamed despite a high bacterial burden. Heavy intestinal colonization with *Salmonella* does not affect chicken weight gain, in agreement with previous work.

We confirmed that the gut microbiota of untreated chicks undergoes the expected dynamic reorganization during first two weeks of chick development [21–23, 27–29] and results in the replacement of the pioneer Proteobacterial population by Firmicutes. Interestingly, Bacteroidetes, reported to be another common phylum in the chick ceca [30–32], was not abundant in the microbial community of the Leghorn chicks we used. Perhaps diet and geographic location, known to play a role in the modulation of chicken microbiota [33, 34], contributed to this difference.

Introduction of STm to the developing microbial community results in a major shift in the phyla in the cecal population. We detected a noticeable increase in Lactobacillales abundance previously linked with the presence of *Salmonella* in the chicken gut [35]. The presence of STm may create a microaerophilic environment in the gut lumen, thereby promoting growth of lactic acid bacteria and limiting the expansion of Clostridia.

The gut microbiome as a complex ecosystem makes an important contribution to chicken metabolism by providing enzymes for the breakdown of macronutrients and the synthesis of vitamins. Although chickens can survive without ceca [36], this organ plays an important role in recycling of urea, water regulation, and carbohydrate fermentation [37–39]. Well-functioning ceca cover approximately 10 percent of chicken energy needs [39]. However, the full fermentation capacity of ceca in broilers is not reached before 28 days of age [40]. Non-targeted analysis of cecal metabolites reflected the dynamic modulation of the microbiota. As expected [41], lactate, the main product of glucose fermentation by *Enterococcus* [42], was detected in ceca of uninfected chicks at low levels. Accordingly, the increased presence of *Enterococcus gallinarum* in the gut of STm infected chicks, resulted in higher accumulation of this metabolite.

Our current understanding of metabolic changes caused by *Salmonella* colonization in the intestine is based on cecum metabolite profile alterations caused by *Salmonella* Enteritidis inoculation of two-week-old UCD-003 layer chicks [43]. Our metabolomic analysis showed a limited overlap with this published data [43]. This discrepancy is likely due to the differences in *Salmonella* serotypes used, and to differences in the ages of the chicks at the time of infection. Nevertheless, at least two previously identified metabolites, isopropylmalic acid and 4-hydroxybenzoic acid, were consistently more abundant in the ceca of chickens infected with either *Salmonella* Enteritidis [43], or *Salmonella* Typhimurium. Isoprolylmalic acid is an intermediate product in the biosynthesis of leucine from 2-keto-valine. Our metagenomic analysis confirmed the increased abundance of branched-chain amino acids biosynthetic genes in the ceca of STm infected chicks mainly originating from two bacterial species, *Salmonella enterica*, and *Enterococcus gallinarum*.

Branched-chain amino acids (BCAA), isoleucine, valine, and leucine, play an important role in the growth, production performance, immunity, and intestinal health of chickens. Inclusion of BCAA in the chicken diet is essential for the proper functioning of the immune system and the maintenance of intestinal mucosal integrity [44]. However, our current understanding of the effect of BCAA on bacterial diversity in the gut is very limited. BCAA are quickly depleted from pig intestines by the microbial community [45]. In turn, supplementation of protein-restricted piglet diet with BCAA facilitated overgrowth of Lactobacillales [46]. BCAA restrictions from the diet resulted in increased susceptibility of mice to *Salmonella* Typhimurium [47]. Thus, BCAA biosynthesis plays an important role in *Salmonella* pathogenesis of orally infected mice [48]. Our data demonstrate that the importance of this biosynthetic pathway not only for survival in mammalian host but also for successful *Salmonella* engraftment in the developing microbial community of young chicks.

Our multifaceted approach reveals the consequences on microbial community structure caused by *Salmonella* colonization, reveals the changes in metabolites present in the altered gut environment and illustrates the requirement for branched-chain amino acid biosynthetic pathway for chicken gut colonization.

## MATERIALS AND METHODS

### Bacterial Strains and Media

Bacterial strains used in this study are listed in Table 1. Salmonella strains used in this study were derived from *Salmonella enterica* subsp. *enterica*, serovar Typhimurium ATCC® 14028s (ATCC, Manassas, VA). Strains were routinely grown in Luria-Bertani (LB) broth or M9 minimal media with appropriate antibiotics. When strains were used for infection of chicks, overnight cultures were grown at 41°C with aeration and appropriate antibiotics. Antibiotics and amino acids were used in the following concentrations: 100 mg/ml nalidixic acid (Nal), 100 mg/ml chloramphenicol, 100 mg/ml carbenicillin (Carb), 500 mg/L L-isoleucine, 500 mg/L L-valine.

A deletion mutant lacking *STM3909 (*HA1632, Δ*STM3909*), the mutant complemented with a wild type copy of *STM3909* (HA1633, Δ*STM3909* bearing *pWSK29::STM3909*) and the wild-type (WT, HA420) strains were grown in Luria-Bertani (LB) broth and in M9 minimal media with or without amino acid supplementation, at 37°C with aeration, and were plated on LB agar plates containing appropriate antibiotics.

### Plasmid construction

A plasmid containing an intact *ilvC* gene (*STM3909*; 1476 bp), with flanking regions ~200 bp upstream the gene, was purchased from GenScript® Biotech. In short, an oligonucleotide encompassing this region was synthesized and cut with KpnI and BamHI restriction enzymes and ligated into pWSK29 cut with the same enzymes. This plasmid was transformed into HA1632 (*ΔSTM3909*) to generate the complemented mutant HA1633 (*ΔSTM3909, pWSK29::STM3909*).

### Ethics Statement

All animal experiments were conducted in compliance with the Guide for the Care and Use of Laboratory Animals of the National Institutes of Health. The protocol has been reviewed and approved by the Animal Care and Use Committee (IACUC) of Texas A&M University (AUP permit 2017-0137).

### Chick hatching

Specific Pathogen Free (SPF) eggs (Charles River SPA-FAS) were incubated in an egg hatchery (GQF Manufacturing Co., Savannah, GA) at 38°C with 58-65% humidity during a 21-day hatching period. During the first 18 days, the eggs were turned 2-3 times a day and were moved to a hatching tray during the last 3 days of the hatching period, where they remained still, to allow the chicks to hatch. Newly hatched chicks were moved to warmed brooders and maintained at 32-35°C. Birds had *ad libitum* access to water and irradiated antibiotic-free 3958 Teklad Laboratory Chick Diet (Envigo Teklad, Cambridgeshire, UK).

### Individual infection in chicks

Seventy-seven chicks were hatched and either left unperturbed (control), or infected with STm HA420. 38 chicks were infected with STm by gavage on day 4 post-hatch with 1 × 10^8^ organisms. Inocula were serially diluted and plated for accurate determination of the concentration of organisms in the inoculum. Animals in both groups were monitored daily for weight change and for signs of infection. At two-day intervals beginning on day 2 and up to day 16 post hatch, 4-5 animals from each group were euthanized and ileal, cecal, colonic contents, as well as liver and spleen were collected for CFU enumeration. A final group of chicks from each treatment group was euthanized at day 19 post hatch. Cecal contents were also used for microbiota analysis and metabolic profiling.

### Competitive infection in chicks

SPF chicks (N = 36) were orally infected with an inoculum of 1×10^8^ CFU of a 1:1 mixture of either WT HA420 and HA1632 (*ΔSTM3909*), or WT (HA420) and HA1633 (*ΔSTM3909 pSTM3909*) on day 4 post-hatch. After infection, chicks were monitored twice daily for signs of disease. On days 2, 8, 12 and 15 post-infection (6,12, 16 and 19 days post-hatch) 4-8 chicks were humanely euthanized. The ceca, ileum, colon, spleen, and liver were excised, homogenized in PBS, and serially diluted and plated on LB plates with appropriate antibiotics for CFU enumeration to calculate the competitive index between the two infecting strains.

### 16S rDNA sequencing

Cecal contents collected from chicks at days 2, 4, 6, 8, 10, 12, 14, 16 and 19 post hatch were flash frozen in liquid nitrogen and stored at −80°C. Bacterial DNA was extracted from cecal content using QIAamp PowerFecal DNA kit (Qiagen). The 16s rRNA V3-4 variable region was amplified with 515F (GTGYCAGCMGCCGCGGTAA) and 806R (GGACTACNVGGGTWTCTAAT) primers to construct Illumina compatible sequencing libraries. All libraries were purified using gel electrophoresis to remove background amplification and primer dimers. Libraries were sequenced to generate 150bp paired-end reads using the Illumina MiSeq platform. The raw 16S rRNA sequence data were processed with QIIME 2 (Quantitative Insights into Microbial Ecology) software package [49]. OUT (operational taxonomic units) picking was done using the Silva database [50].

### Shotgun metagenomic sequencing

Bacterial DNA was extracted from the cecal contents collected at day 4, 6, 10, 12 and 19 post-hatch from uninfected at day 6, 10, 12 and 19 post-hatch from *Salmonella*-infected chicks as described above and quantified using Qubit dsDNA fluorometric assay (Thermo Fisher). DNA quality was assessed by DNA ScreenTape assay using TapeStation 2200 (Agilent). Samples were normalized and used to generate sequencing libraries using the Nextera DNAflex Library preparation kit following the manufacturer’s protocol (Illumina). Normalized libraries were pooled in equimolar ratio, and then sequenced on a single lane of a NovaSeq S4 2×150 flow cell (NovaSeq 6000, Illumina) with at least 50 million reads per sample.

### Data processing

After sequencing, a total of approximately 2.3 billion paired end 150bp reads were generated across thirty-six samples. The reads were then processed using bioBakery whole genome sequencing (WGS) metagenomic pipeline [51]. Raw reads were processed using Kneaddata (https://huttenhower.sph.harvard.edu/kneaddata/), to remove adapter sequences and host (*Gallus gallus*) reads. Specifically, Kneaddata workflow included trimming of overrepresented reads with FastQC (https://qubeshub.org/resources/fastqc) followed by removal of adapter sequences using Trimmomatic [52]. The remaining reads were further filtered using Bowtie 2 tool [53] to remove chicken related host reads. Filtered reads were used as an input for taxonomic profiling and strain identification using MetaPhlAn2 [54] and StrainPhlAn2 [55], respectively. HUMAnN2 tool [56] was then used for functional profiling and for detection of abundant pathways in the sample-specific pangenomes. Unmapped reads remaining after this step were used to perform a translated search against the Uniref90 database using DIAMOND. Collected results were used for quantification of the relative pathway abundances.

### Gas Chromatography Mass Spectroscopy metabolic profiling

Cecal contents of infected and uninfected birds were collected, resuspended in sterile PBS, weighed, and placed on ice. Samples were vortexed for 2 minutes and then separated by centrifugation at 6000 x g for 15 minutes at 4°C. Supernatants were mixed with 5 μM of deuterated lactate (sodium L-lactate-3,3,3,-d_3_, CDN Isotopes) as an internal standard, transferred to the new tubes and dried using a SpeedVac concentrator. Samples were dissolved by addition of pyridine in 1:1 ratio, sonicated for 1 minute and incubated at 80°C for 20 minutes. Next, the derivatization reagent *N-tert*-Butyldimethylsilyl-*N*-methyltrifluoroacetamide with 1 % t-BDMCS (*tert*-Butyldimethylchlorosilane, Cerilliant) was added and samples were incubated at 80 °C for 1 hour. After centrifugation at 14,000 rpm for 1 minute, derivatized samples were transferred to autosampler vials for gas chromatography- mass spectrometry (GC-MS) analysis (Shimadzu, TQ8040) using 30 m × 0.25 mm × 0.25 μm Rtx-5Sil MS column (Shimadzu). The injection temperature was 250°C and the injection split ratio was set to 1:100 with an injection volume of 1 μL. The oven temperature started at 50°C for 2 minutes, increasing to 100°C at 20°C per minute and to 330°C at 40°C per minute with a final hold at this temperature for 3 minutes. Flow rate of the helium carrier gas was kept constant at a linear velocity of 50 cm/s. The interface temperature was 300°C. The electron impact ion source temperature was 200°C, with 70 V ionization voltage and 150 μA current. For qualitative experiments, Q3 scans (range of 50-550 m/z, 1000 m/z per second) were performed, and putative compounds were identified by searching the Shimadzu database.

To quantitatively measure lactate and deuterated lactate, multiple reaction monitoring was used with target ion m/z 261>233 and a reference ion m/z 261>189 for lactate, and target ion m/z 264>236 and a reference ion m/z 264>189 for deuterated lactate, respectively.

---

**Figure S1. Non-targeted metabolomics analysis of the chicken cecal content.** Cecal content was collected from uninfected (left panel) and STM-infected (right panel) chicks and analyzed by GC-MS.

## REFERENCES

1. Thorns, C., Bacterial Food-borne zoonoses. Rev Sci Tech, 2000. 19(1): p. 226–39.

2. Rodrigue, D., R. Tauxe, and B. Rowe, International increase in Salmonella enteritidis: a new pandemic. Epidemiol. Infect, 1990. 105: p. 21–27.

3. Majowicz, S., et al., The global burden of nontyphoidal Salmonella gastroenteritis. Clin. Infect. Dis., 2010. 50: p. 882–9.

4. CDC, National enteric disease surveillance: salmonella annual report, 2014. 2014, Centers for Disease Control and Prevention: Atlanta, GA.

5. CDC, Antibiotic Resistance Threats in the United States, in U.S. Department of Health and Human Services. 2019, Centers for Disease Control and Prevention: Atlanta, GA.

6. Van Immerseel, F., et al., Intermittent long-term shedding and induction of carrier birds after infection of chickens early posthatch with a low or high dose of Salmonella Enteritidis. Poult Sci, 2004. 83: p. 1911–1916.

7. Smith, H. and J. Tucker, The effect of antibiotic therapy on the faecal excretion of Salmonella typhimurium by experimentally infected chickens. J. Hyg. (London), 1975. 75: p. 275–292.

8. Beal, R., et al., Age at primary infection with Salmonella enterica serovar Typhimurium in the chicken influences persistence of infection and subsequent immunity to re-challenge. Vet Immunol Immunop, 2004. 100: p. 151–164.

9. Nacomcf, F., Response to Questions Posed by the Food Safety and Inspection Service Regarding Salmonella Control Strategies in Poultry. J Food Prot, 2019. 82(4): p. 645–668.

10. Wigley, P., Salmonella enterica in the chicken: how it has helped our understanding of immunology in a non-biomedical model species. Front Immunol, 2014. 5(482).

11. Spiga, L., et al., An Oxidative Central Metabolism Enables Salmonella to Utilize Microbiota-Derived Succinate. Cell Host Microbe, 2017. 22(3): p. 291–301.

12. Stecher, B., et al., Salmonella enterica serovar Typhimurium exploits inflammation to compete with the intestinal microbiota. PLoS Biol, 2007. 5(10): p. e244.

13. Behnsen, J., et al., Exploiting host immunity: the Salmonella paradigm. Trends Immunol, 2015. 36(2): p. 112–120.

14. Khan, C., The Dynamic Interactions between Salmonella and the Microbiota, within the Challenging Niche of the Gastrointestinal Tract. International Scholarly Research Notices, 2014. 2014: p. 846049.

15. Rhen, M., Salmonella and Reactive Oxygen Species: A Love-Hate Relationship. Journal of innate immunity, 2019. 11(3): p. 216–226.

16. Faber, F., et al., Respiration of Microbiota-Derived 1,2-propanediol Drives Salmonella Expansion during Colitis. PLoS pathog, 2017. 13(1): p. e1006129.

17. Eade, C., et al., Salmonella Pathogenicity Island 1 Is Expressed in the Chicken Intestine and Promotes Bacterial Proliferation. Infection and immunity. Infect Immun, 2018. 87(1): p. e00503–18.

18. Gillis, C.C., et al., Dysbiosis-Associated Change in Host Metabolism Generates Lactate to Support Salmonella Growth. Cell Host Microbe, 2018. 23(1): p. 54–64 e6.

19. Winter, S.E., C.A. Lopez, and A.J. Baumler, The dynamics of gut-associated microbial communities during inflammation. EMBO Rep, 2013. 14(4): p. 319–27.

20. Apajalahti, J.K., A.; Graham, H., Characteristics of the gastrointestinal microbial communities, with special reference to the chicken. World’s Poultry Science Journal 2004. 60(2): p. 223–232.

21. Tanikawa, T., et al., Aging transition of the bacterial community structure in the chick ceca. Poult Sci, 2011. 90(5): p. 1004–8.

22. Awad, W.A., et al., Age-Related Differences in the Luminal and Mucosa-Associated Gut Microbiome of Broiler Chickens and Shifts Associated with Campylobacter jejuni Infection. Front Cell Infect Microbiol, 2016. 6: p. 154.

23. Ballou, A.L., et al., Development of the Chick Microbiome: How Early Exposure Influences Future Microbial Diversity. Front Vet Sci, 2016. 3: p. 2.

24. Chasser, K.M., et al., Enteric permeability and inflammation associated with day of hatch Enterobacteriaceae inoculation. Poult Sci, 2021. 100(9): p. 101298.

25. Ghareeb, K.A., W.A.; Bohm, J.; Zebeli, Q., Impact of luminal and systemic endotoxin exposure on gut function, immune response and performance of chickens. World’s Poultry Science Journal, 2016. 72(2): p. 367–380.

26. Kogut, M.H. and R.J. Arsenault, Immunometabolic Phenotype Alterations Associated with the Induction of Disease Tolerance and Persistent Asymptomatic Infection of Salmonella in the Chicken Intestine. Front Immunol, 2017. 8: p. 372.

27. Wilson, K.M., et al., Evaluation of the impact of in ovo administered bacteria on microbiome of chicks through 10 days of age. Poult Sci, 2019. 98(11): p. 5949–5960.

28. Litvak, Y., et al., Commensal Enterobacteriaceae Protect against Salmonella Colonization through Oxygen Competition. Cell Host Microbe, 2019. 25(1): p. 128–139 e5.

29. van der Wielen, P.W., et al., Spatial and temporal variation of the intestinal bacterial community in commercially raised broiler chickens during growth. Microb Ecol, 2002. 44(3): p. 286–93.

30. Wei, S., M. Morrison, and Z. Yu, Bacterial census of poultry intestinal microbiome. Poult Sci, 2013. 92(3): p. 671–83.

31. Oakley, B.B., et al., The chicken gastrointestinal microbiome. FEMS Microbiol Lett, 2014. 360(2): p. 100–12.

32. Sergeant, M.J., et al., Extensive microbial and functional diversity within the chicken cecal microbiome. PLoS One, 2014. 9(3): p. e91941.

33. Pin Viso, N., et al., Geography as non-genetic modulation factor of chicken cecal microbiota. PLoS One, 2021. 16(1): p. e0244724.

34. Biasato, I., et al., Modulation of intestinal microbiota, morphology and mucin composition by dietary insect meal inclusion in free-range chickens. BMC Vet Res, 2018. 14(1): p. 383.

35. Videnska, P., et al., Influence of Salmonella enterica serovar Enteritidis infection on the composition of chicken cecal microbiota. BMC Vet Res, 2013. 9: p. 140.

36. Chaplin, S.B., Effect of cecectomy on water and nutrient absorption of birds. J Exp Zool Suppl, 1989. 3: p. 81–6.

37. Karasawa, Y., Significant role of the nitrogen recycling system through the ceca occurs in protein-depleted chickens. J Exp Zool, 1999. 283(4-5): p. 418–25.

38. Bjornhag, G., Transport of water and food particles through the avian ceca and colon. J Exp Zool Suppl, 1989. 3: p. 32–7.

39. Jozefiak, D.R., A.; Martin, S.A., Carbohydrate fermentation in the avian ceca: a review. Animal Feed Science and Technology, 2004. 113: p. 1–15.

40. Svihus, B.C., M.; Classen, H.L., Function and nutritional roles of the avian caeca: a review. World’s Poultry Science Journal, 2013. 69(2): p. 249–264.

41. Jozefiak, D., et al., The effect of beta-glucanase supplementation of barley- and oat-based diets on growth performance and fermentation in broiler chicken gastrointestinal tract. Br Poult Sci, 2006. 47(1): p. 57–64.

42. Hatti-Kaul, R., et al., Lactic acid bacteria: from starter cultures to producers of chemicals. FEMS Microbiol Lett, 2018. 365(20).

43. Mon, K.K.Z., et al., Integrative analysis of gut microbiome and metabolites revealed novel mechanisms of intestinal Salmonella carriage in chicken. Sci Rep, 2020. 10(1): p. 4809.

44. Kim, W.K., et al., Functional role of branched chain amino acids in poultry: a review. Poult Sci, 2022. 101(5): p. 101715.

45. Dai, Z.L., et al., Utilization of amino acids by bacteria from the pig small intestine. Amino Acids, 2010. 39(5): p. 1201–15.

46. Yin, J., et al., Branched-chain amino acids, especially of leucine and valine, mediate the protein restricted response in a piglet model. Food Funct, 2020. 11(2): p. 1304–1311.

47. Petro, T.M. and J.K. Bhattacharjee, Effect of dietary essential amino acid limitations upon the susceptibility to Salmonella typhimurium and the effect upon humoral and cellular immune responses in mice. Infect Immun, 1981. 32(1): p. 251–9.

48. Fitzsimmons, L.F., et al., Salmonella Reprograms Nucleotide Metabolism in Its Adaptation to Nitrosative Stress. mBio, 2018. 9(1).

49. Bolyen, E., et al., Reproducible, interactive, scalable and extensible microbiome data science using QIIME 2. Nat Biotechnol, 2019. 37(8): p. 852–857.

50. Quast, C., et al., The SILVA ribosomal RNA gene database project: improved data processing and web-based tools. Nucleic Acids Res, 2013. 41(Database issue): p. D590–6.

51. McIver, L.J., et al., bioBakery: a meta’omic analysis environment. Bioinformatics, 2018. 34(7): p. 1235–1237.

52. Bolger, A.M., M. Lohse, and B. Usadel, Trimmomatic: a flexible trimmer for Illumina sequence data. Bioinformatics, 2014. 30(15): p. 2114–20.

53. Langmead, B. and S.L. Salzberg, Fast gapped-read alignment with Bowtie 2. Nat Methods, 2012. 9(4): p. 357–9.

54. Truong, D.T., et al., MetaPhlAn2 for enhanced metagenomic taxonomic profiling. Nat Methods, 2015. 12(10): p. 902–3.

55. Truong, D.T., et al., Microbial strain-level population structure and genetic diversity from metagenomes. Genome Res, 2017. 27(4): p. 626–638.

56. Franzosa, E.A., et al., Species-level functional profiling of metagenomes and metatranscriptomes. Nat Methods, 2018. 15(11): p. 962–968.

